# Huntingtin is essential for synaptic plasticity in the adult hippocampus

**DOI:** 10.1101/2022.10.05.510980

**Authors:** Jessica C. Barron, Firoozeh Nafar, Matthew P. Parsons

## Abstract

Huntingtin (HTT), an exceptionally large protein with hundreds of interacting partners within the central nervous system, has been extensively studied due to its role in Huntington’s disease (HD) pathology. HD is a monogenic disorder caused by a polyglutamine repeat expansion in the *HTT* gene, which results in the production of a pathogenic mutant huntingtin (mHTT) protein, and toxic effects of this mutant protein in the context of HD have been well-established. Less-established, however, is the role of wild type HTT (wtHTT) in the adult brain, particularly in areas outside the corticostriatal pathway. wtHTT has previously been suggested to play a vital role in cellular functions that promote synapse homeostasis, such as fast axonal transport of synaptic cargo, vesicle replenishment and receptor localization and stability. Synaptic dysfunction precedes and predicts cell death in many neurodegenerative diseases including HD (termed synaptopathies) and whether proper synaptic transmission can be maintained without wtHTT in extrastriatal brain areas such as the hippocampus remains unknown. Consequences of wtHTT reduction in the adult brain are of particular importance as clinical trials for many non-selective HTT-lowering therapies for HD are underway, which are unable to distinguish between mHTT and wtHTT, and therefore reduce levels of both proteins. We investigated the consequences of wtHTT loss of function in the CA3-CA1 pathway of the adult hippocampus using a conditional knockout mouse model and found that 1-2 month deletion of wtHTT in excitatory hippocampal neurons inhibits post-tetanic potentiation and completely abolishes NMDA receptor-dependent long-term potentiation in these animals. These data reveal a novel role of wtHTT as an essential regulator of short- and long-term plasticity in the adult hippocampus.

## Introduction

The huntingtin (HTT) protein is a structurally large protein that is ubiquitously expressed in the body but has particular relevance to the central nervous system due to its role in Huntington’s disease (HD), an autosomal dominant neurodegenerative disease caused by a CAG repeat expansion in the *HTT* gene. Within the brain, wild type HTT (wtHTT) was found by initial studies to have a few hundred interacting partners (reviewed in Harjes and Wanker, 2003; Li and Li, 2004), however, over 700 candidate proteins were later identified in a comprehensive study that incorporated samples from various brain regions that interact with non-mutated HTT including the striatum, cortex and cerebellum (Shirasaki et al., 2012). It is therefore unsurprising that wtHTT has been found to play a role in various fundamental cellular processes including fast axonal transport, transcription, autophagy and cell survival signalling (Saudou and Humbert, 2016). In addition, due to the fact that many of its protein interactors support synapse homeostasis, wtHTT has been suggested as an essential regulator of synaptic function (Barron et al., 2021). However, the role of wtHTT in synaptic transmission, particularly at synapses outside the corticostriatal pathway, such as the hippocampus, remains unclear. Understanding the consequences of wtHTT reduction at hippocampal synapses is vital to understanding HD, as synaptic plasticity impairments in the Schaffer collateral pathway of the hippocampus have been shown to correlate with cognitive dysfunction in HD mouse models (Milnerwood et al., 2006). Cognitive symptoms are reported by caregivers of HD patients to be the most burdensome aspects of the disease and are known to manifest over a decade earlier than motor symptoms (Paulsen, 2011). Furthermore, premanifest HD patients show impairments in hippocampal-based cognitive tasks, which positively correlate to number of years to symptom onset (Begeti et al., 2016). Impairments in long term potentiation (LTP) at Schaffer collateral synapses in symptomatic HD rodent models have been well-established (Hodgson et al., 1999; Lynch et al., 2007; Murphy et al., 2000; Quirion and Parsons, 2019; Usdin et al., 1999). As well, our lab has recently shown that presymptomatic HD mice, in addition to hippocampal LTP dysregulation, also show activity-dependant alterations in their pre and postsynaptic nanostructures, revealed by super-resolution analysis (Ravalia et al., 2021). Whether wtHTT is essential for LTP facilitation and maintenance at the CA3-CA1 synapse, however, remains unknown. Understanding the consequences of wtHTT loss in the CA3-CA1 pathway is essential, as many HD genetic therapies are well underway, which aim to lower total HTT protein levels in the nervous system. Of these therapies, antisense oligonucleotides (ASOs) have been some of the most promising drug candidates thus far with many advancing past Phase 1 clinical trials (Tabrizi et al., 2022, 2019). However, many of these ASOs are non-selective in that they cannot distinguish between mutated HTT and wtHTT RNA, and therefore will lower levels of both proteins throughout the adult brain, including in extrastriatal structures such as the hippocampus. It is clear that wtHTT is essential for nervous system development (Duyao et al., 1995; Nasir et al., 1995; Zeitlin et al., 1995), however, the consequences of wtHTT depletion in the adult mammalian brain using therapeutics such as ASOs, particularly in areas outside the corticostriatal pathway remain unknown and additional research at the preclinical level is required immediately to understand how this may impact health outcomes following non-selective HTT lowering.

wtHTT is known to play a prominent role in the bidirectional transport of synaptic cargo by forming a complex with its interactor HAP1 and molecular motors kinesin and dynein (Engelender, 1997; Li et al., 1998; McGuire et al., 2006). In addition to promoting anterograde and retrograde transport of the neurotrophin BDNF (Gauthier et al., 2004), it has also been shown to regulate bidirectional transport of its tyrosine kinase receptor, TrkB (Liot et al., 2013). BDNF has been shown to be essential for LTP induction and maintenance in the hippocampus, and retrograde trafficking of BDNF-bound TrkB to the cell soma receptors promotes survival signalling (Zheng et al., 2008). wtHTT may also be involved in ensuring delivery of other important neurotrophins that support synaptic plasticity, as wtHTT loss has been shown to disrupt the transport of dense core vesicles, which specialize in the transport of neurotrophins and neuropeptides (Bulgari et al., 2017). Moreover, wtHTT is also a positive regulator of BDNF at the transcriptional level (Zuccato et al., 2003, 2001).

Given the suggested role of wtHTT as a major regulator of synaptic plasticity and its prominent role in BDNF transport and transcription, we investigated the consequences wtHTT protein deletion in the adult mouse hippocampus by conditionally deleting wtHTT in excitatory hippocampal neurons. We found that wtHTT cKO animals had significantly reduced post-tetanic potentiation as well as impaired LTP in comparison to control animals after 1-2 months of protein deletion. Data shown here support our hypothesis that wtHTT is essential for synaptic plasticity in the adult mammalian hippocampus.

## Materials and Methods

### Animals

Mice used for experiments were initially delivered to our institution with permission from Dr. Scott Zeitlin’s laboratory (Dragatsis et al., 2000). These colonies were then maintained in Memorial University of Newfoundland’s Animal Care Services facility. Mice were biologically engineered to have their endogenous wild type huntingtin alleles flanked with loxP sites in order to conditionally inactivate the mouse *Htt* gene in a homozygous manner by Cre injection (*Htt*^fl/fl^). Mice were bred as homozygous x homozygous. Both male and female mice were used for all experiments. Mice were group housed in ventilated cage racks and kept on a 12 h light/dark cycle (lights on at 7:00 A.M.) with food and water available *ad libitum*. All experimental procedures were approved by Memorial University’s Institutional Animal Care Committee and were performed in accordance with the guidelines set by the Canadian Council on Animal Care.

### Stereotaxic Surgery

*Htt*^fl/fl^ mice aged 2-4 months were anesthetized using isoflurane (3% induction, 1.5–2% maintenance) and injected subcutaneously in the abdominal region with 2 mg/kg meloxicam and below the scalp with 0.1 ml/0.2% lidocaine. Above the desired brain coordinates two small holes were drilled in the skull using an Ideal Micro-Drill (Harvard Apparatus). A Model 7002 KH Neuros Hamilton Syringe was used along with an infusion pump (Pump 11 Elite Nanomite, Harvard Apparatus) to inject 1 μl pENN.AAV.CamKII.HI.GFP-Cre.WPRE.SV40 (Addgene, catalog # 105551-AAV9) or 1 μl pENN.AAV.CamKII0.4.eGFP.WPRE.rBG (Addgene, catalog #105541-AAV9) bilaterally into the dorsal hippocampus (injection rate, 2 nl/s). The following coordinates were used with respect to distance from bregma: 2.6 mm posterior, 2.3 mm medial/lateral and 1.1–1.3 mm ventral to brain surface. The syringe was left in place for at least 5 min following the injection before careful removal from the brain. 0.5 ml of 0.9% saline was injected subcutaneously in the abdominal region and the scalp incision was sutured. Mice were monitored for 20-30 minutes as they recovered post-surgery on a heating pad before being returned to their designated home cage area. All experiments were performed 1-2 months following stereotaxic surgery, unless otherwise stated.

### Acute Slice Preparation

Mice were anesthetized using isoflurane before decapitation and whole brain removal. The brain was then immersed in ice-cold oxygenated (95% O2/5% CO2) slicing solution consisting of (in mM): 125 NaCl, 2.5 KCI, 25 NaHCO3, 1.25 NaH2PO4, 2.5 MgCl2, 0.5 CaCl2, and 10 glucose. A Precisionary compresstome was used to obtain transverse acute hippocampal slides from both hippocampi. Slices were transferred to a holding chamber containing oxygenated ACSF to recover for at least 1.5 hours before experimentation. ACSF used during slice recovery and electrophysiological recordings consisted of (in mM) 125 NaCl, 2.5 KCI, 25 NaHCO3, 1.25 NaH2PO4, 1 MgCl2, 2 CaCl2, and 10 glucose.

### Multi-electrode Array Electrophysiology

All electrophysiological experiments were performed using a MED64 multi-array system (Alpha MED Scientific) using a protocol previously described by our lab (Ravalia et al., 2021). Slices were positioned on the array for stimulation and subsequent recording of synapses within the Schaffer collateral pathway. Mobius software (Alpha Med Scientific) was used to select the optimal CA3 stimulating electrode based on its ability to evoke a clear, stable fEPSP response in CA1 stratum radiatum. Data from one CA1 recording electrode per slice was selected for analysis, also based on these criteria. An input-output curve was recorded by stimulating between 10 μA and 120 μA in 10μA increments, and baseline fEPSPs were evoked using a stimulation intensity that resulted in ∼35% of the maximal response. A stable baseline was acquired for at least 10 minutes before beginning the LTP stimulation protocol, which was induced by one train of theta burst stimulation (TBS) consisting of 10 bursts with each burst comprised of 4 pulses at 100 hz (200 ms inter-burst interval). Recordings were held for 50 minutes following LTP induction. Before the LTP experiment, paired pulse facilitation was measured in each slice by evoking two fEPSPs separated by 50, 100 or 250 ms.

### Western Blotting

For Western blot quantification of HTT protein expression, dorsal hippocampi of AAV-Cre-GFP or AAV-eGFP-injected *Htt*^fl/fl^ mice were dissected and homogenized in 400 ul of Lysis buffer containing Halt Protease and Phosphatase inhibitor Cocktails (Thermo Scientific, catalog #78440). The supernatant was collected, and the protein concentration was determined using BCA standards. Fifty micrograms of protein was added to each lane of a Bolt 4-12% Bis-Tris Plus gel for electrophoresis before being transferred to 0.45 um nitrocellulose membrane. Primary antibodies for HTT (1:1000; mouse-monoclonal, Sigma Aldrich catalog #MAB2166) and beta-actin (1:5000; mouse-monocolonal, Sigma Aldrich catalog #A5316 clone AC74), and a goat anti-mouse IgG secondary antibody (1:5000; monoclonal; Thermofisher Scientific catalog #31430) were used. Blots were developed using a chemiluminescent HRP substrate (Super Signal West Pico Plus Thermo Scientific catalog #34580). Band densities were quantified in ImageJ, and HTT band densities were normalized to actin.

### Statistics

All statistical tests were performed using version 9.4.1 GraphPad Prism and included unpaired t-tests and two-way repeated-measures ANOVA followed with the appropriate post-hoc tests. Specific statistical tests used are described in results text. P values less than 0.05 were considered significant.

## Results

### Conditional Knockout of the Huntingtin Protein for 1-2 Months Impairs Short- and Long-Term Plasticity at CA3-CA1 Hippocampal Synapses

Bioengineered *Htt*^fl/fl^ mice received bilateral hippocampal injections of CaMKII-promoted Cre-recombinase viral vector AAV-GFP-Cre specifically targeting dorsal CA1 at 2-4 months of age to conditionally KO (cKO) wtHTT in excitatory hippocampal neurons of adult mice. Control animals were injected with CaMKII-promoted AAV-eGFP using the same hippocampal coordinates. Clear GFP expression was detected in the pyramidal layer in CA1 (Fig 1A). In our hands, huntingtin antibodies are more reliable with western blot compared to immunohistochemistry; therefore, wtHTT protein levels in control and cKO mice were quantified by western blotting from dorsal hippocampal samples collected 1-2 months post-AAV injection. Cre injections resulted in clear wtHTT reduction, as cKO animals showed a 55.7% decrease in wtHTT protein (Fig. 1B, n=8 hemispheres from 4 animals per condition. Unpaired t-test. p <0.0001). As Cre was expressed only in excitatory neurons via the CaMKII promoter, complete loss of wtHTT is not observed with wester blot, and the remaining wtHTT is likely to represent its unaltered expression in inhibitory interneurons and glia.

**Figure 1.**
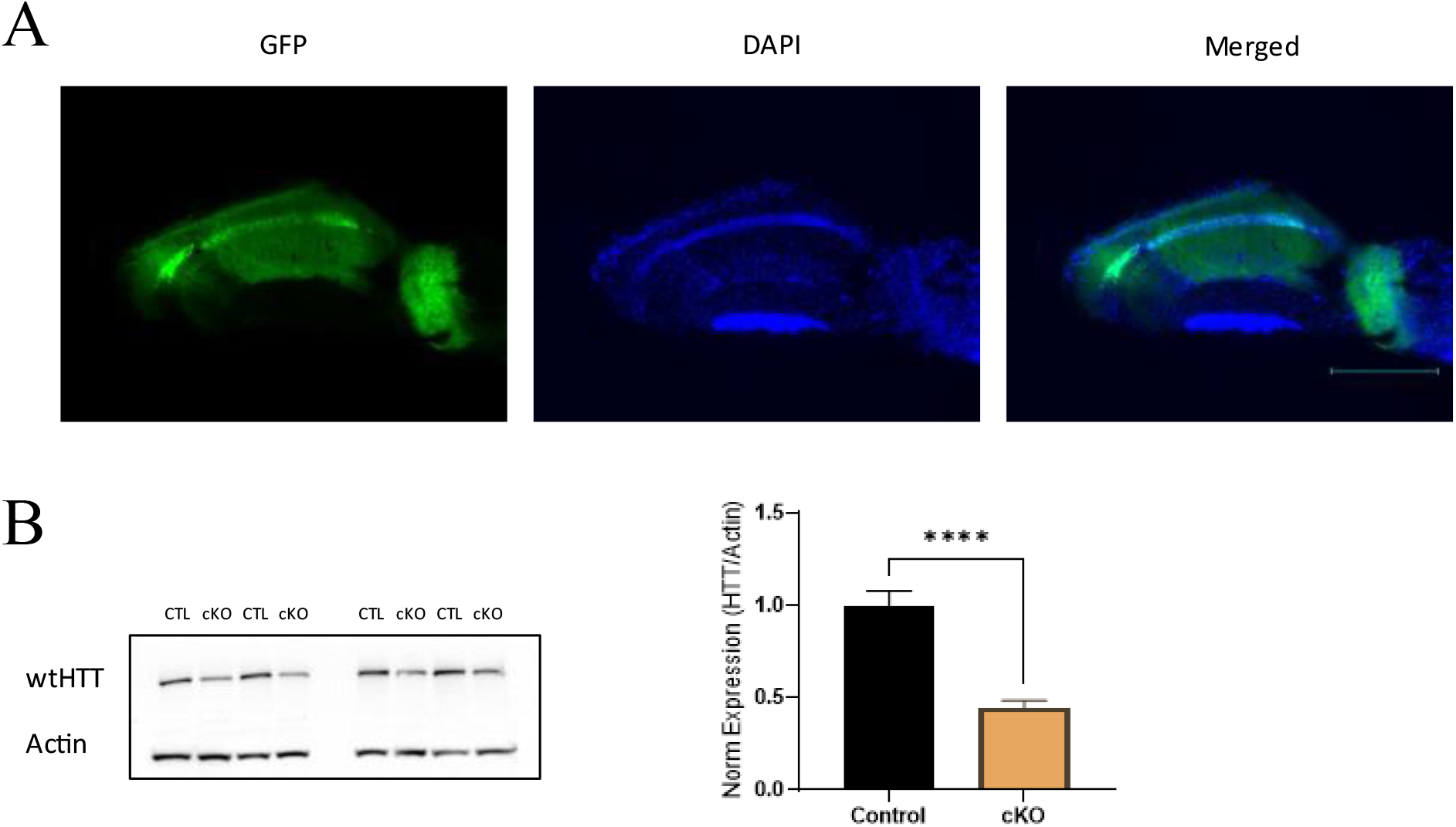
wtHTT is conditionally deleted in excitatory cells of the CA3-CA1 hippocampi of *Htt*fl/fl mice using Cre recombinase system. (A) Representative images showing GFP expression and DAPI-stained nuclei in acute hippocampal slice from control mouse injected with CaMKII-AAV-eGFP bilaterally into the dorsal hippocampus. Image acquired 1-2 months post-injection. Scale bar signifies 750 μm (B) Western blotting results showing 55.7% reduction in wtHTT from dorsal hippocampal tissues samples of control (CTL) and wtHTT cKO mice. Unpaired t-test. n=8 hemispheres from 4 animals (2 males, 2 females) per condition. Error bars indicate standard error of the mean. **** p < 0.0001.

In separate mice, electrophysiological experiments were performed on acute hippocampal slices to measure synaptic plasticity using a MED64 multielectrode array system. GFP expression was used to confirm accurate AAV targeting to the hippocampus. After achieving a stable baseline, LTP was induced by 1 train of TBS (10 bursts each consisting of 4 pulses at 100 hz with 200 ms between bursts). As expected, TBS increased in the slope of the fEPSP measured 45-50 minutes after LTP induction in control mice (Fig. 2A, n=8-11 acute slices, n=6-7 animals per condition). In contrast, in cKO mice, fEPSP responses returned to baseline levels within just a few minutes following TBS, and LTP was completely abolished (Fig. 2B, unpaired t-test. p < 0.05). In addition to the striking LTP impairment in cKO mice, wtHTT loss in adulthood also significantly inhibited post-tetanic potentiation (PTP), measured 0-2 minutes after TBS (Fig. 2C, unpaired t-test. p < 0.01). Basal synaptic transmission, measured using standard input-output curves (Fig 3A), and paired-pulse ratios did not differ between wtHTT cKO mice in comparison to control animals (Fig 3B), although paired-pulse facilitation tended to be a bit lower in cKO mice compared to controls (two-way ANOVA. CTL vs cKO p=0.061).

**Figure 2.**
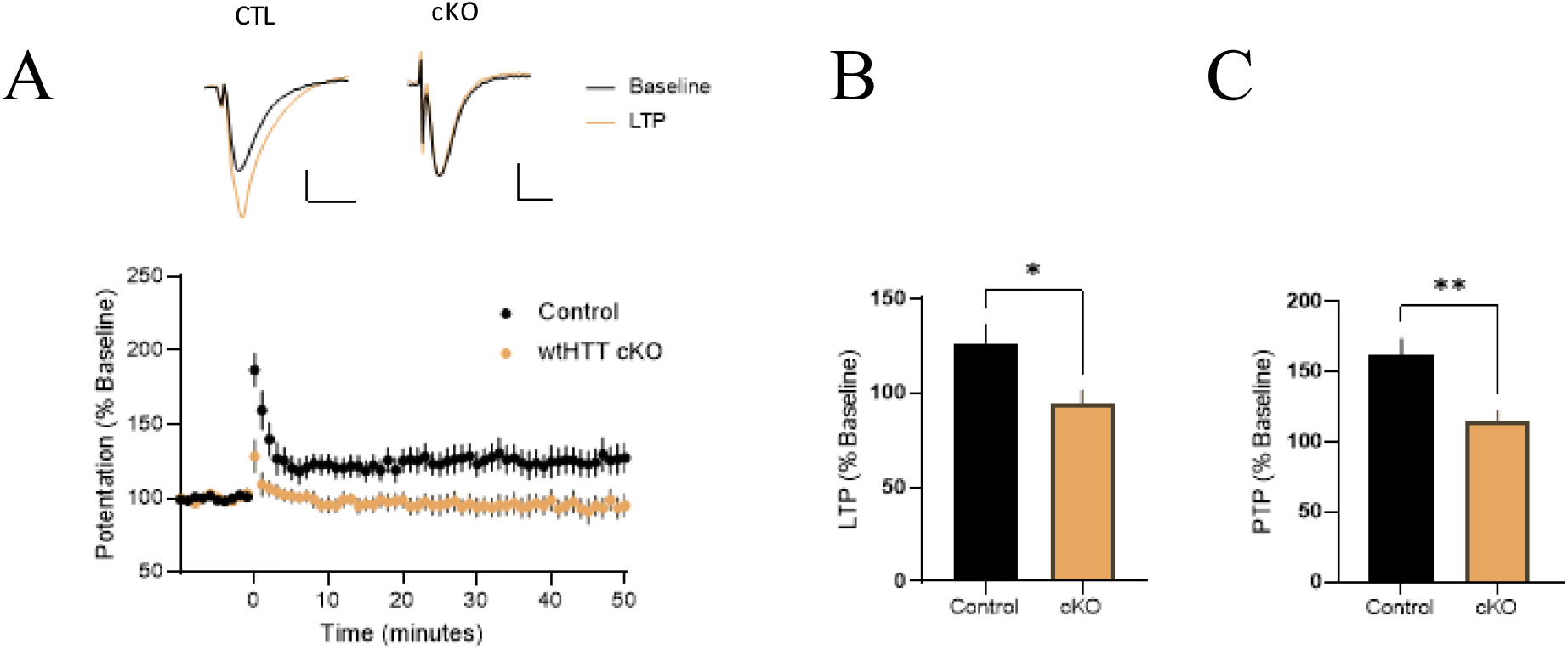
1-2 month conditional knockout of wtHTT in CA3-CA1 excitatory neurons results in short- and long-term plasticity deficits. (A) Control and cKO LTP recordings after 1 train of TBS stimulation (10 bursts consisting of 4 pulses at 100 hz with 200 ms between bursts) at t=0 minutes, shown as percentage compared to baseline recorded for at least 10 minutes before stimulation. n=8-11 acute slices from 6-7 animals per condition. CTL representative trace axes: 0.1 mV, 20 ms. cKO representative trace axes: 0.05 mV, 20ms. (B) LTP (long-term potentiation) measured 45-50 minutes after TBS. Control animals show a 26.6% increase while wtHTT cKO mice show a 5.16% decrease compared to baseline fEPSP recording. (C) Post-tetanic potentiation (PTP) measured 0-2 minutes after TBS. Control animals show a 62.1% increase while cKO mice show a 14.8% increase. Unpaired t-tests. Error bars indicate standard error of the mean. *p < 0.05, ** p < 0.01.

**Figure 3.**
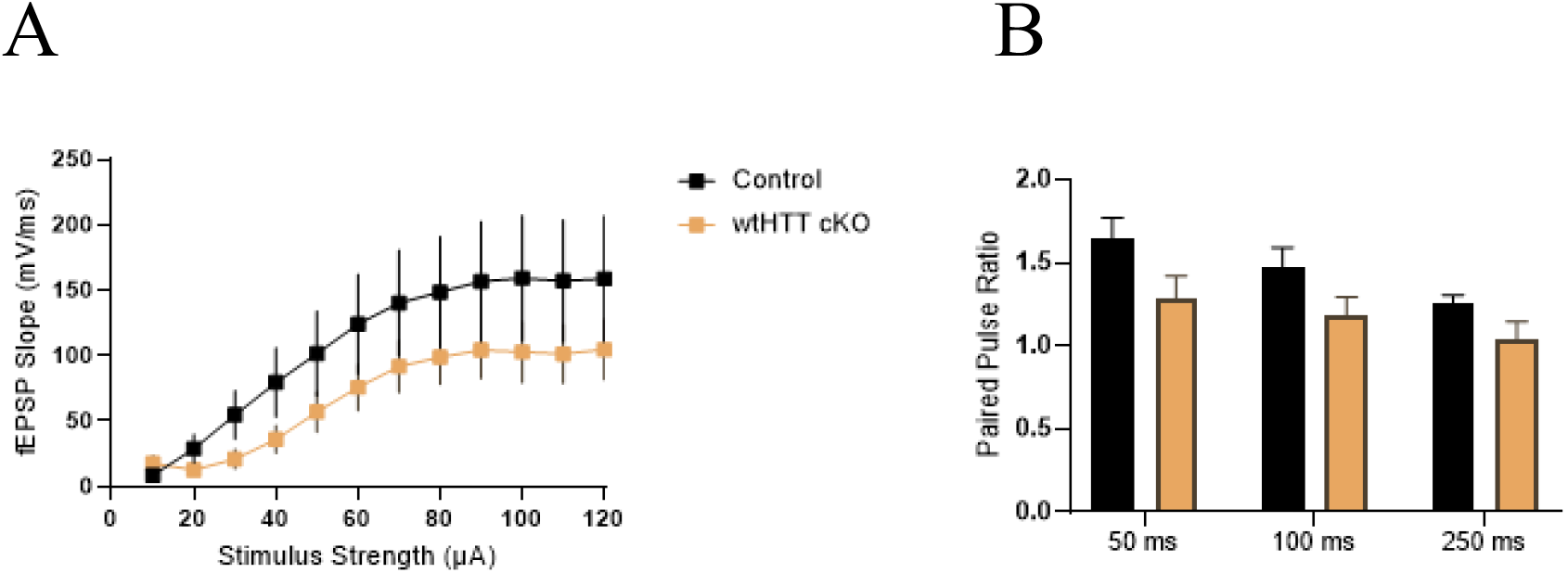
1-2 month wtHTT conditional knockout in CA3-CA1 excitatory cells does not significantly impair basal synaptic transmission or paired pulse facilitation in cKO mice. (A) Standard input output curve recorded by stimulating from 10 μA to 120 μA in 10μA increments and recording slope of subsequent field excitatory postsynaptic potentials (fEPSPs). Two-way ANOVA comparing CTL vs cKO treatment shows p=0.323. n=8-11 acute slices, n=6-7 animals per condition. (B) Paired pulse ratios were determined by evoking two baseline fEPSPs separated by 50, 100 and 250 milliseconds. Two-way ANOVA comparing CTL vs cKO treatment shows p=0.061. n=8-11 acute slices, n=6-7 animals per condition.

### Long-Term Knockout of the wtHTT Protein for 6-8 Months Abolishes fEPSPs at Schaffer collateral Synapses

We found that knocking out wtHTT in excitatory neurons in the adult hippocampus severely impairs PTP and LTP 1-2 months later. As non-selective HTT lowering aims to reduce HTT levels for a longer period of time, we also asked whether sustained wtHTT knockout would result in a similar effect, or if additional mechanisms could eventually compensate for the loss of wtHTT function. To this end, we injected a cohort of control and cKO mice to be used for electrophysiological experiments 6-8 months after AAV injection. GFP presence confirmed the long-term expression of transgenes. Unexpectedly, while we able to generate clear fEPSP responses and typical LTP experiments in these older control animals, we were unable to evoke clear fEPSP responses from cKO animals; therefore, CA1 recordings of LTP were not even possible at this stage in our hands. This observation suggests that sustained wtHTT loss negatively impacts basal connectivity at CA3-CA1 synapses. Together, our findings demonstrate that wtHTT loss of function in adulthood can have striking negative consequences on the basal and activity-dependent properties of hippocampal synapses.

## Discussion

Due to its extensive number of protein interactors, there are numerous ways in which wtHTT may regulate hippocampal plasticity, aside from its well-established role as positive regulator of BDNF protein levels and its intracellular transport. For instance, wtHTT increases AMPA receptor transport to the cortical postsynapse through its complex with HAP1 and kinesin motor KIF5 (Mandal et al., 2011), and wtHTT-lowering decreases clustering of the postsynaptic scaffolding protein PSD-95 in mixed corticostriatal cultures (Parsons et al., 2014). This combination of less available AMPA receptors to insert into postsynaptic slots and instability of PSD-95 in the postsynaptic density could lead to lower postsynaptic sodium influx upon glutamate release and/or fewer available AMPA receptors to insert into the postsynaptic membrane upon NMDA receptor activation post-TBS, which could reflect the loss of LTP seen in our wtHTT cKO mice. Additionally, wtHTT affects the palmitoylation of various proteins through its associations with the palmitoyl acetyltransferases HIP14 and HIP14L. Particularly, HIP14L has been shown to palmitoylate cluster II of the GluN2B NMDA receptor subunit in striatal neurons, which increases the presence of GluN2B-containing receptors at the postsynaptic membrane (Kang et al., 2019). NMDA receptor composition is especially relevant to HD, as extrasynaptic GluN2B-containing receptors are associated with cell-death signalling pathways in HD mouse models (Lundh et al., 1991). wtHTT increases the enzymatic activity of its interactor HIP14 (Huang et al., 2011) and it has been suggested that it may also increase the palmitoylation activity of HIP14L, based on the structural similarities between HIP14 and HIP14L. Thus, wtHTT loss may impact synaptic function and plasticity by decreasing the palmitoylation of key synaptic proteins.

wtHTT has been previously shown to influence the structural stability of spines. Embryonic cKO of wtHTT in mice has been shown to alter synaptic density postnatally in cortical neurons. Interestingly, an increase in synapse number and accelerated spine maturation was seen after the first three weeks of life, however, this excited state could not be maintained as 5 week old cKO cortical neurons (where Htt in Htt^(flox/-)^ mice was inactivated by Emx1-Cre) have fewer spines and a more immature profile (McKinstry et al., 2014). As well, it was recently shown that when wtHTT is depleted embryonically in excitatory cortical neurons, postsynaptic cells specifically have altered spine morphology alongside a significant increase in the presence of mature mushroom spines and a decrease in immature filapodia, driven by changes in the RAC1/LIMK/cofilin signaling pathway (Wennagel et al., 2022). Both studies also found that these structural changes correlated with decreased cortical synaptic activity, based on patch-clamp electrophysiological recordings. Deletion of wtHTT from distinct subpopulations of dopamine neurons in the striatum also results in alterations in synapse density, where wtHTT cKO mice showed a decrease in inhibitory synapses from indirect pathway SPNs whereas an increase in inhibitory synapses was seen in direct pathway SPNs (Burrus et al., 2020).

In all, results shown here in conjunction with our previous review of the role of wtHTT at the synapse (Barron et al., 2021), give substantial evidence for wtHTT as a major regulator of synaptic transmission and plasticity. These data suggest that non-selective HTT-lowering in the adult brain may lead to failures in pathways that underlie hippocampal-dependant learning and memory if wtHTT is depleted in this important brain area. Given that non-selective HTT lowering therapeutics are in clinical trials, it is of interest for future studies to identify the precise mechanisms by which wtHTT loss in adulthood impairs synaptic plasticity, and to determine how much wtHTT loss, if any, can be tolerated at the synaptic level.

## Acknowledgements

All research presented in the current manuscript was funded by Dr. Parsons’ Canadian Institutes of Health Research (CIHR) project grant. We would like to thank the Animal Care Services staff at Memorial University for their assistance with the housing, monitoring and care of all research animals used for these experiments.

